# A Stickiness Response System in Rats

**DOI:** 10.64898/2026.08.06.742745

**Authors:** Sandra Tan, Simone Rencken, Tom Childs, Jasmine T. Stone, Adam J. Tiesman, Pamela N. Anderson, Michael Brecht, Ann M. Clemens

## Abstract

The perception of stickiness is known to everyone who interacts with the world. While eating, walking, and navigating diverse environments including crowded subways, forests, fields and lunchrooms, stickiness is a common and old sensation. Responses to sticky stimuli have been measured in animal and human brains; however, precise behavioral responses and the underlying neural mechanisms are not well understood. We applied sticky stimuli to three-week-old rat pups and found the effects vary greatly across the animal’s body: Sticky stimuli are quickly removed from forepaws and nose, but often evoke only little reaction from hindpaws. When we applied sticky (marshmallow, mochi) and non-sticky stimuli (water, oil) to forepaws, we observed stimulus unspecific behaviors (licking and grooming) with variable response onsets as well as three fast-onset sticky-specific behaviors. Sticky-specific behaviors were exclusively triggered by sticky stimuli and included paw shaking and paw swiping (behaviors presumably aiming at stickiness removal) and paw tapping. In tapping, animals gently tap their forepaws onto each other or on the ground; we wondered if the resulting paw-substrate detachments serve stickiness sensing. Blocking of forepaw skin sensation reduced sticky-specific responses to sticky stimuli compared to control conditions (Ringer’s injections). To assess central representations of stickiness, we obtained *in vivo* whole-cell recordings of neurons in forepaw-somatosensory-cortex while presenting sticky and non-sticky stimuli to anesthetized rat pups. While responses were heterogeneous across the population, we observed individual neurons that had significantly different responses to stimulus detachment for sticky and non-sticky stimuli. In summary, we describe a fast-onset, body-part-specific stickiness response system in rats, which is strongly driven by forepaw skin afferents.

**Graphical abstract:** 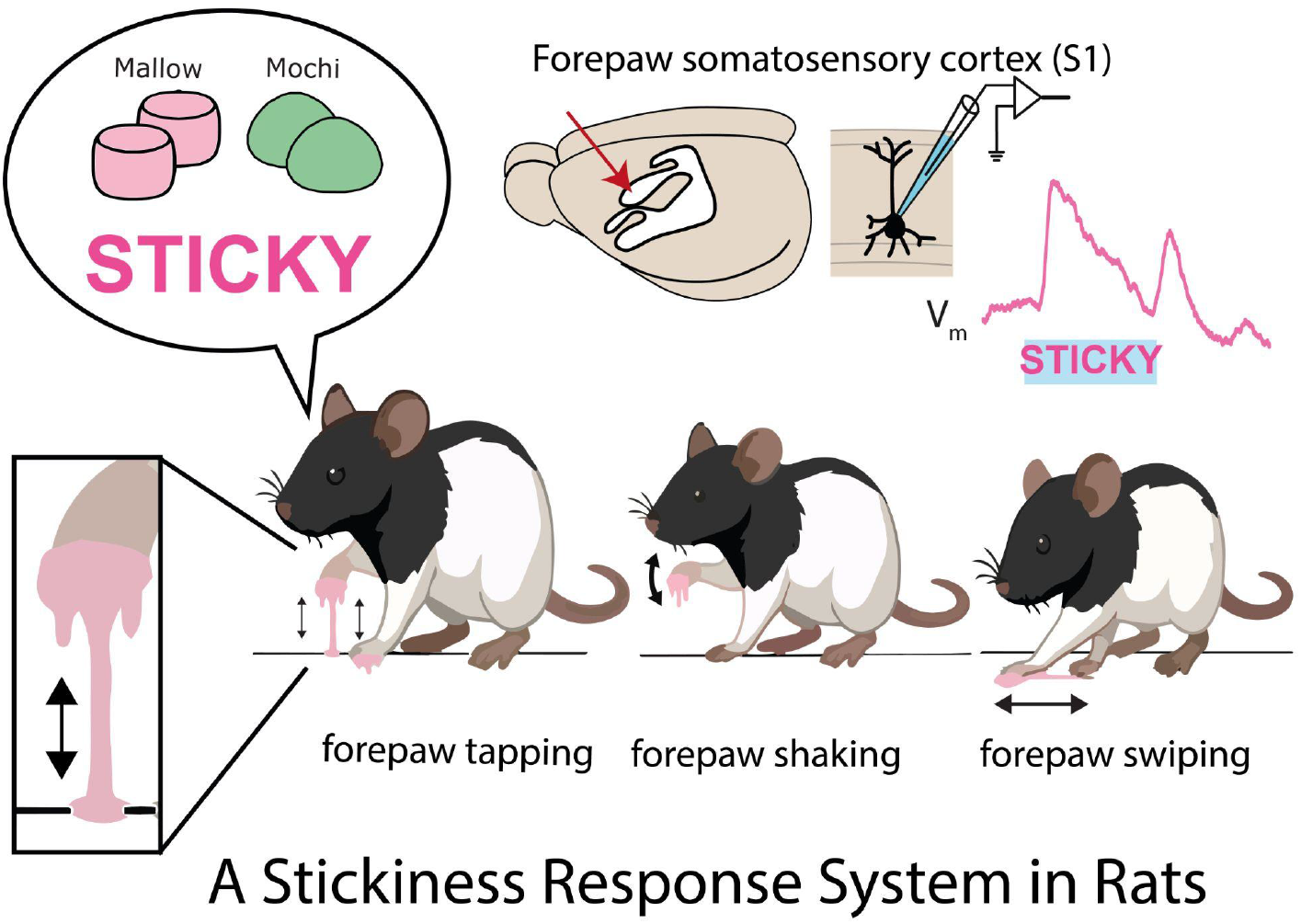

## Introduction

Stickiness is an everyday phenomenon sensed immediately and vividly. Everyone is familiar with the stickiness of stimuli such as honey. While we most commonly experience stickiness in the tactile domain, the mechanisms for stickiness perception are remarkably broad and elaborate. Accordingly, we infer stickiness from the rhythmic sounds of a sticky attachment on the soles of our shoes even in the absence of direct touch. Languages onomatopoeically mimic the sticky sounds of food^1^. Given the sophistication of everyday stickiness perception, we are remarkably ignorant about the underlying neural mechanisms.

Judging tactile sensations by subjective introspection, stickiness perception appears to be quite distinct from other tactile qualities such sharpness, roughness, smoothness and the like. Specifically, we perceive the latter qualities when contacting tactile surfaces; stickiness, in contrast, emerges, when we detach from surfaces and experience the pull of sticky stimuli.

The analysis of stickiness by rigorous quantitative analysis is a relatively young endeavor. Nam et al. (2020) studied psychophysically how stickiness judgments vary with physical characteristics of surface detachment from the fingertip ^2^. This work confirmed the intuition that detachment forces shape stickiness perception.

Pioneering work on the rodent whisker system related the stickiness of surfaces to vibrissa kinematics during vibrissal touch^3^. Other investigators studied more derived qualities of sticky stimuli, namely their ability to evoke tactile disgust ^4,5^.

Another branch of investigation determined brain regions activated by sticky stimuli in the human brain ^6,7^. Yeon et al. 2017 described sticky-stimulus-related activity in the human somatosensory cortex and dorsolateral prefrontal cortex. In follow-up experiments these investigators also described multisensory integration mechanisms for stickiness perception ^8^.

We sought to combine behavioral and neural approaches to understand stickiness at a single-neuron level. We did so in rats and investigated their behavioral and neural responses to sticky stimuli. Specifically, we asked the following questions: (i) Do rats react to sticky stimuli and if yes, how do such responses depend on the body location to which sticky stimuli are applied? (ii) Are specific behaviors evoked by sticky stimuli? (iii) What is the temporal organization of stickiness-evoked behaviors? (iv) Do stickiness-evoked behaviors depend on tactile sensations? (v) Do somatosensory cortical neurons respond to and differentiate between sticky and non-sticky stimuli?

## Results

### Behavioral responses to sticky stimuli across the body

To quantitatively assess behavioural responses to stickiness, we applied sticky stimuli across the body (**Figure 1A**). We observed removal behaviors after application of marshmallow to the nose, ventral forepaw, dorsal forepaw, nape, ventral hindpaw, and dorsal hindpaw in 600-second trials (n=6 animals) (**Figure 1B**). The time to first foregroom behavior was variable across body location (Friedman test, Q = 21.88, p = 0.0006). Time to first removal attempt also showed significant differences across body locations (Friedman test, Q = 25.39, p = 0.0001). For both measures of removal behaviors the nose was the quickest location (mean proportion of total time: first foregroom = 0.011, first removal attempt = 0.011), while the hindpaw and nape remained the slowest with several rats failing to initiate any removal behaviors from the ventral hindpaw within the 600s window (mean proportion of total time = 1.00 for first removal attempt) (**Figure 1C**).

**Figure 1:**
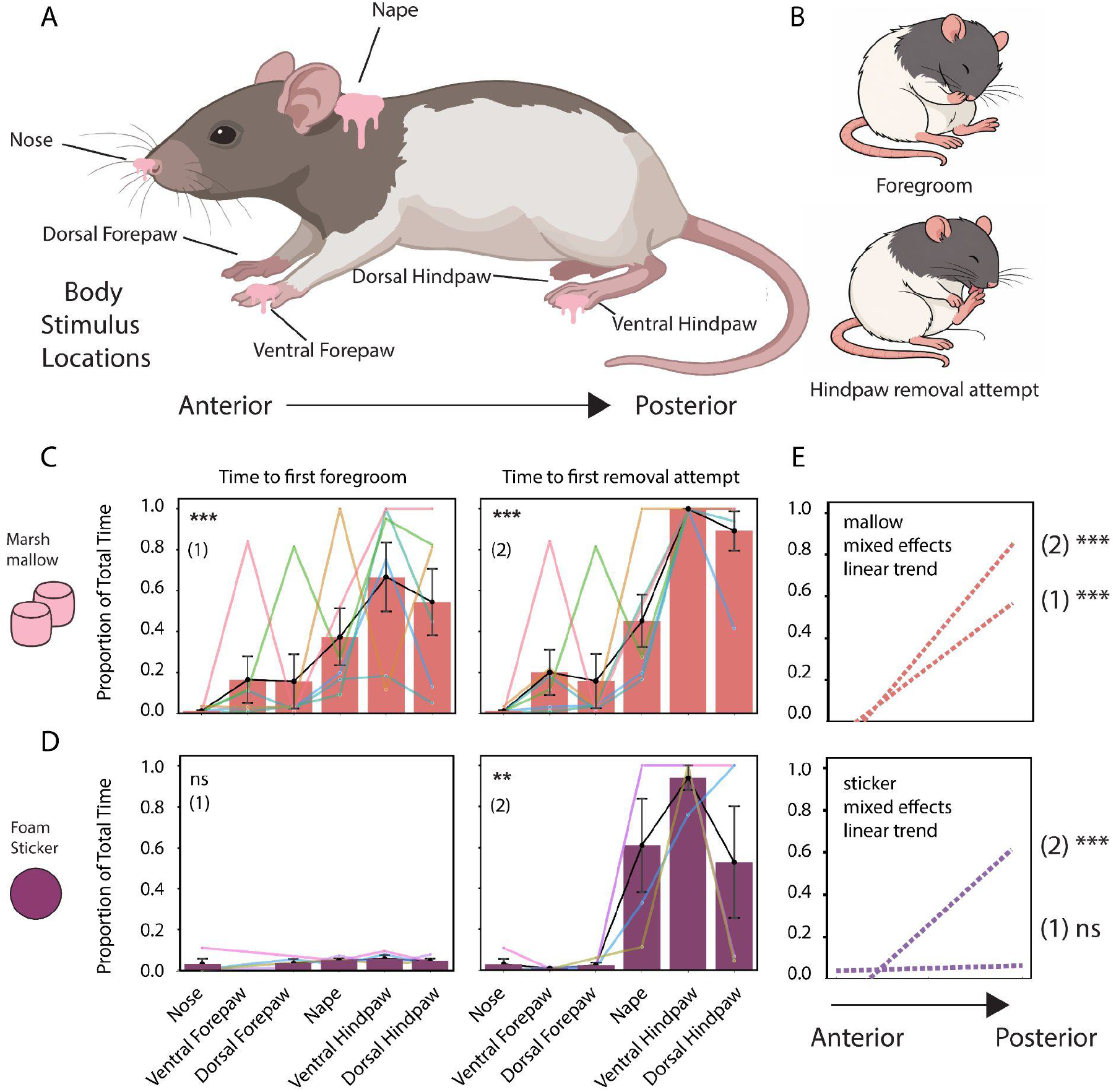
Behavioral responses to sticky stimuli across the body. **A**) Schematic of the locations where the stimulus was applied. In the anterior rat body, stimulus was applied to the nose, ventral forepaw, and dorsal forepaw. In the posterior rat body, stimulus was applied to the nape, ventral hindpaw, and dorsal hindpaw. **B)** Depiction examples of scored behaviors after stimulus application. Time to first foregroom (top) and first removal attempt (hindpaw), following application of stimuli to bodyparts. **C)** Marshmallow was applied to each body part in separate 600 sec trials. Barplots show mean ± SEM, and the black line represents the mean across individuals. Non-black points represent individual rats with non-black lines connecting across body locations for that individual (n = 6). Trials in which the behavior did not occur within 600s were assigned a value of 1. Left: Barplot showing the proportion of total time taken to first foregroom after application of marshmallow across body locations (Friedman test, Q = 21.88, p < 0.0006). Right: Barplot showing the proportion of total time taken to the first removal attempt of the marshmallow across body locations (Friedman test, Q = 25.39, p = 0.0001). **D)** A foam sticker was applied to each body part in separate 600 sec trials. Barplots show mean ± SEM, and the black line represents the mean across individuals. Non-black points represent individual rats with non-black lines connecting across body locations for that individual (n = 4). Trials in which the behavior did not occur within 600s were assigned a value of 1. Left: Barplot showing the proportion of time taken to first foregroom after application of foam sticker across body locations (Freidman test, Q = 4.80, p =0.308). Right: Barplot showing the proportion of time taken to first removal attempt of the foam sticker across body locations (Friedman test, Q = 15.60, p = 0.0081). **E)** Anterior-to-posterior slopes across body locations show significant positive slope across both measures for marshmallow (mixed-effects linear trend: p= 2.96e-08; p= 3.00e-19). For the foam sticker, a non-significant trend was seen for first foregroom (mixed-effects linear trend: ns, p = 0.204) and positive slope trends were seen for first removal attempt and time to successful removal (mixed-effects linear trend: p = 2.71e-06). Detailed slope analyses and additional information shown in **Figure S1**. * p<0.05, ** p<0.01, *** p<0.001.

We used a foam sticker as the sticky stimulus and saw similar results (n = 4). The first time to foregroom behavior did not differ statistically across body locations (Friedman test, Q = 4.80, p =0.308); note that Ventral Forepaw was excluded from this analysis due to trials in which rats directly proceeded to removal without an observable foregrooming response. The proportion of time to show the first removal attempt of the foam sticker showed statistically significant differences across body locations (Friedman test, Q = 15.60, p = 0.0081). The proportion of time to the successful removal of the foam sticker was different across body locations (Friedman test, Q = 17.81, p = 0.0032).

As with marshmallow, the nose showed the fastest response across all three measures (mean proportion of total time: first foregroom = 0.028, first removal attempt = 0.028, successful removal = 0.032), while the hindpaw and nape were slower, with most rats failing to attempt or successfully remove the sticker at these sites (mean proportion of total time > 0.87 for first removal attempt and successful removal at dorsal hindpaw, nape, and ventral hindpaw) (**Figure 1D and Figure S1**).

Post-hoc tests did not show significant pairwise differences between individual body locations for either adhesive after Holm correction for multiple comparisons. However, we also performed a mixed effects linear trend comparing the behavior across the anterior to posterior axis. This analysis showed that anterior body parts were significantly faster across both measures in the marshmallow application (mixed-effects linear trend: p= 2.96e-08; p= 3.00e-19) (**Figure 1E, top and Figure S1**). The same analysis performed on the foam sticker revealed a non-significant trend for the proportion of time to the first foregroom across anterior and posterior body parts (mixed-effects linear trend: ns, p= 0.204), but significant trends for first removal attempt and successful removal (p = 2.71e-06; p = 2.30e-09), consistent with the results from the marshmallow trials, with faster first removal attempts in the anterior body parts compared to the posterior body parts (**Figure 1E, bottom and Figure S1**).

### Stickiness-specific behaviors and their temporal organization

Application of marshmallow to the dorsal forepaw of rat pups evoked a set of stereotyped behaviors that we had not previously observed in a large number of other behavioral contexts. Rats shook the affected forepaw, or both forepaws together (**Figure 2A**; forepaw shaking, **<u>Video 1</u>**); swiped the left and right forepaws across the floor in an alternating fashion (**Figure 2B**; forepaw swiping, **<u>Video 2</u>**); or briefly lifted and replaced the forepaws against the floor, alternating between the left and right paw in quick succession (**Figure 2C**; forepaw tapping, **<u>Video 3</u>**). We define these as ’sticky behaviors’ (**Figure 2A–C**, **<u>Video 4</u>**). Rats also performed bouts of stereotyped self-grooming as well as licking of the adhesive and their forepaws. To analyse the temporal organization of sticky behaviors, we examined their timing across a 180 second trial period following application of marshmallow to the dorsal forepaw (n = 5 rats). We observed that sticky behaviors were performed rapidly after application of the marshmallow and that this led to removal of most of the substance, after which the frequency of sticky behaviors was reduced. In contrast, licking and grooming behaviors continued throughout the 180 second trial period (**Figure 2D**). For each rat and category we measured the half-rise time, defined as the time at which 50% of total observed events were performed (**Figure 2E**). Half-rise times differed across categories (Friedman test, Q = 8.40, p = 0.008), and were earliest for sticky behaviors (19.0 ± 4.8 s), intermediate for licking (42.4 ± 10.4 s), and latest for grooming (69.8 ± 11.5 s; mean ± SEM). Sticky behaviors were the first category to reach half-rise in every rat (5/5). We conclude rats respond to forepaw marshmallow-treatment with fast-onset stereotyped ‘sticky behaviors’ that occur faster than routine body care behaviors such as grooming and licking.

**Figure 2:**
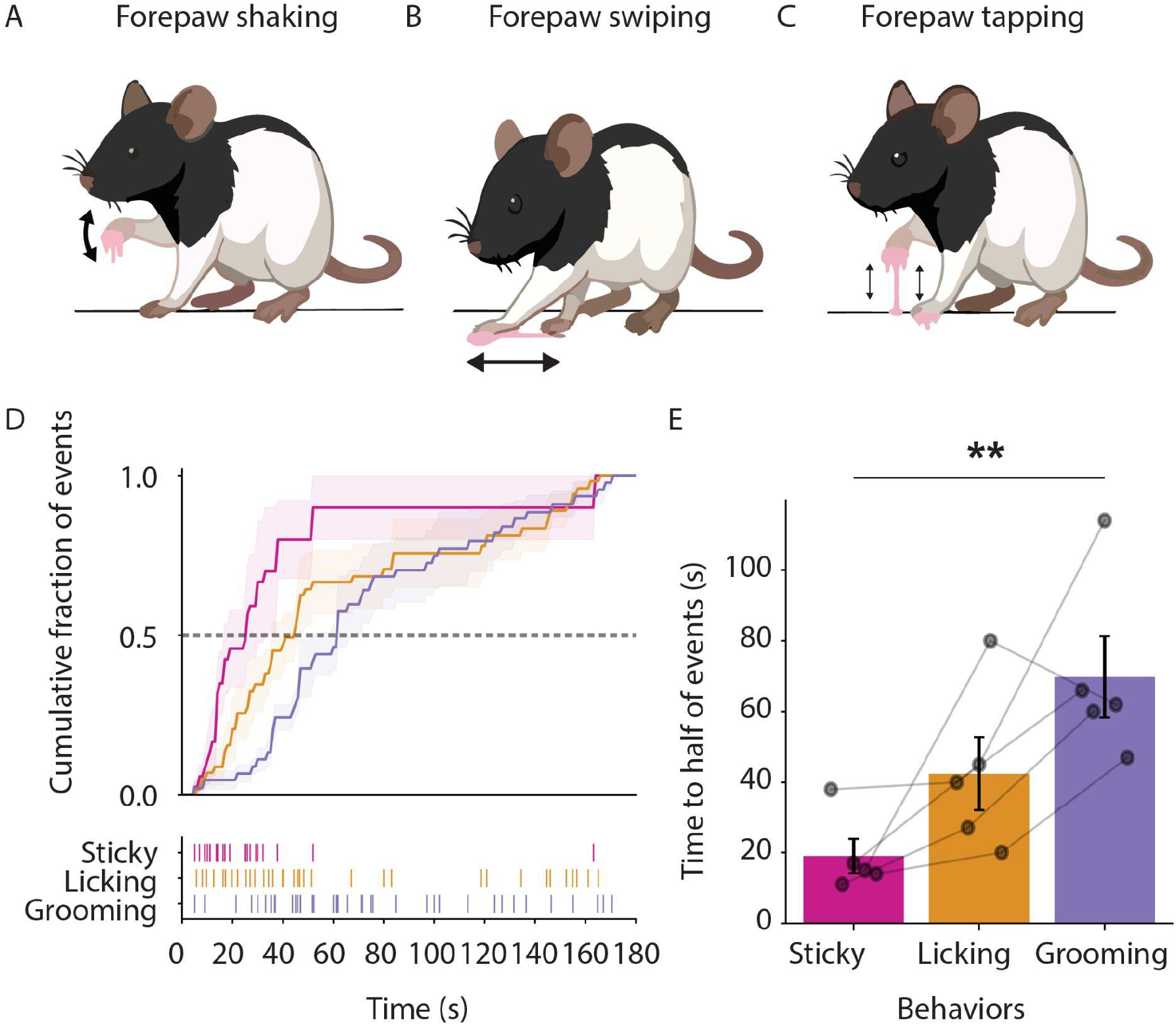
**Stickiness-specific behaviors evoked by marshmallow application to rat forepaw. A–C**) Schematics of three ‘sticky behaviors’. Forepaw shaking (**A, <u>Video 1</u>**) consisted of rapid oscillatory movements of an elevated forepaw, or of both forepaws together; forepaw swiping (**B, <u>Video 2</u>**) consisted of alternating sweeps of the left and right forepaws across the floor; and forepaw tapping (**C, <u>Video 3</u>**) consisted of brief, repeated lifting and replacement of the forepaws against the floor. Pink indicates marshmallow adhesive and arrows indicate movement direction (all behaviors together, **<u>Video 4</u>**). **D**) Top: Normalised cumulative distributions of sticky behaviors (magenta; shaking, swiping, and tapping pooled), licking (orange), and grooming (purple) during the 180 second observation period following application of marshmallow to the dorsal forepaw of P21 Long Evans rats. Cumulative event counts were normalised to the total number of events produced by each rat in each category. Lines and shaded regions show mean ± SEM of the normalised curves (n = 5 rats). Bottom: Raster plot of sticky behaviors, licking, and grooming showing raw event times pooled across the five trials from five rats. **E**) Half-rise times, defined for each rat and behavioral category as the time at which the normalised event count reached 0.5. Bars show mean ± SEM, points represent individual rats, and gray lines connect measurements from the same rat (n = 5). Half-rise times differed significantly among behavioral categories (Friedman test, Q = 8.40, p = 0.008, Kendall’s W = 0.84). Sticky behaviors showed the earliest half-rise times. ** p<0.01

### Sticky-specific behaviors (paw-shaking, -swiping, -tapping) are highly stimulus-specific

To verify that ‘sticky behaviors’ were in fact specific to sticky stimuli we examined rats’ behavioral responses to various sticky and non-sticky stimuli. We applied one of four stimuli—oil, water, mochi, or marshmallow—to each rats’ dorsal forepaw in randomized order (**Figure 3A**). Oil and water served as non-sticky controls whereas mochi and marshmallow served as sticky stimuli. Both grooming (**Figure 3B**) and licking (**Figure 3C**) behaviors were evoked – to varying extents – by all four stimuli applied. This suggests that both grooming and licking are general body cleaning responses. In contrast, sticky behaviors were exclusively evoked by sticky stimuli (mochi and mallow; likelihood-ratio test across stimuli, p=0.0004, Holm-Bonferroni corrected paired comparisons: Oil-water, p=1; Mochi-water, p=0.0009; Mallow-water, p=0.0003; Mochi-oil, p=0.0009; Mallow-oil, p=0.0003; Mallow-mochi, p=1, n=6 rats; **Figure 3D**). We conclude that sticky behaviors are highly stimulus specific.

**Figure 3:**
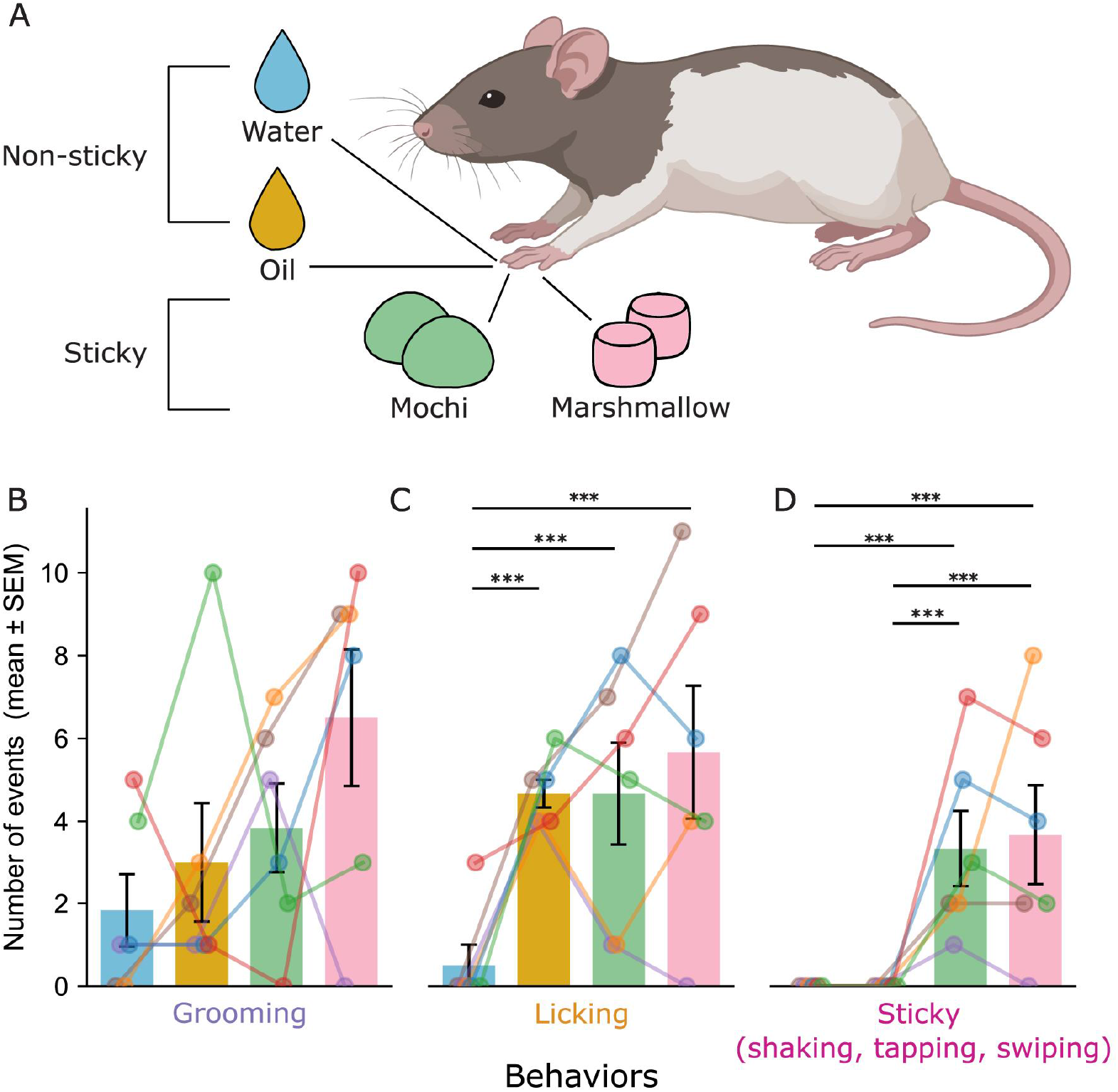
Stereotyped ’sticky’ behaviors occur specifically after application of sticky stimuli. **A)** Schematic of sticky (marshmallow, mochi) and non-sticky (oil, water) stimuli applied to dorsal forepaw. **B-D)** Barplots showing number of removal behaviors (mean ± SEM) after application of various stimuli to the dorsal forepaw. Sticky behaviors were not observed after water (blue) and oil (yellow) application, but were observed after mochi (green) and marshmallow (pink) application. **B)** Grooming stats, mixed effects model, likelihood-ratio test, p=0.065. **C)** Licking mixed effects model, likelihood-ratio test, p=0.001, Holm-Bonferroni corrected paired comparisons: Oil-water, p=0.0009; Mochi-water, p=0.0009; Mallow-water, p=2.34e-05; Mochi-oil, p=1; Mallow-oil, p=1; Mallow-mochi, p=1. **D)** Sticky mixed effects model, likelihood-ratio test, p=0.0004, Holm-Bonferroni corrected paired comparisons: Oil-water, p=1; Mochi-water, p=0.0009; Mallow-water, p=0.0003; Mochi-oil, p=0.0009; Mallow-oil, p=0.0003; Mallow-mochi, p=1. N=6 rats. Bar plots are mean ± SEM. * p<0.05, ** p<0.01, *** p<0.001

### Stickiness-evoked behaviors depend on peripheral tactile sensation

To test whether tactile sensation in the forepaw is important for the execution of sticky behaviors, we injected the local anaesthetic lidocaine into the dorsal forepaw before applying marshmallow and observing removal behaviors (**Figure 4A**). Compared to control injections with Ringer’s solution only (Ringer, n = 6 rats; lidocaine, n = 5 rats), the occurrence of sticky behaviors (paw shaking, swiping, and tapping) was significantly reduced, whereas other removal behaviors such as grooming and licking were unaffected (**Figure 4B**). Data were normally distributed (Shapiro–Wilk test), and two-way ANOVA revealed a significant main effect of treatment (F = 16.31, p = 0.0087). Tukey’s multiple comparisons confirmed a significant reduction in sticky behaviors (adjusted p = 0.0017), with no significant effect on grooming (adjusted p = 0.383) or licking (adjusted p = 0.6019).

**Figure 4:**
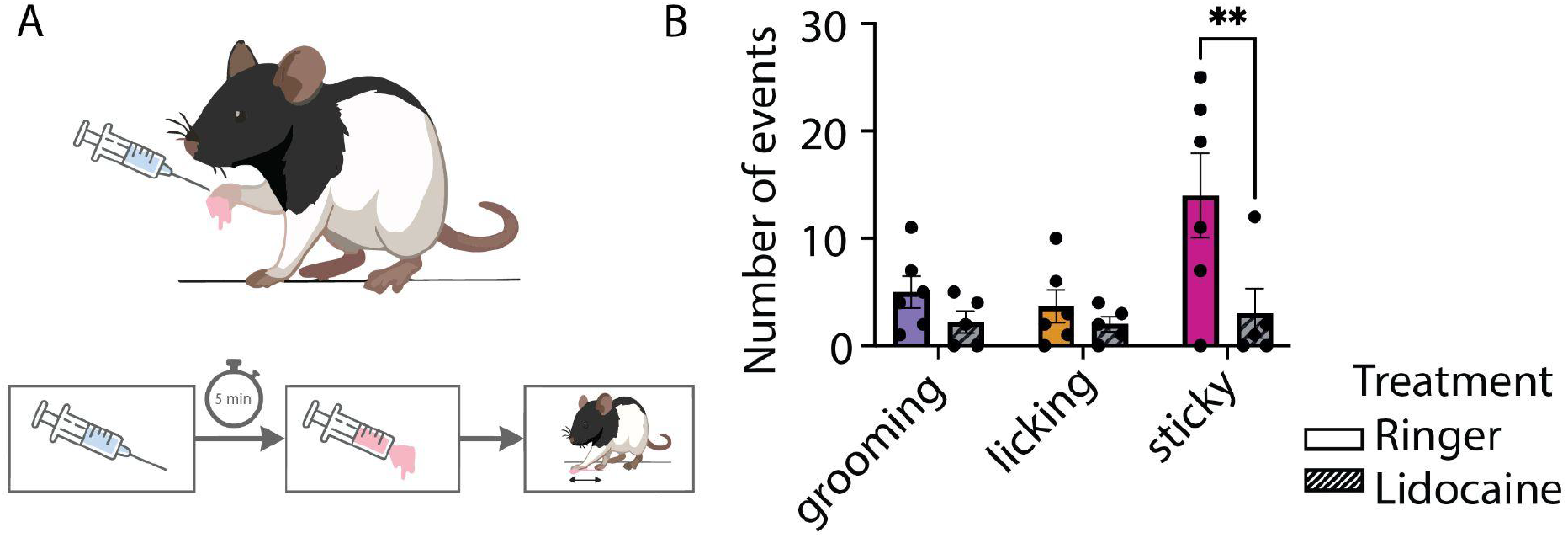
Blocking touch with a local anaesthetic reduces removal behaviors following application of sticky substance. **A)** Experimental overview. Dorsal forepaw of juvenile rats were injected with 0.05 mL of a local anaesthetic (0.5% lidocaine, N = 5) or control (Ringer, N = 6). After 5 min, marshmallow was applied to the injected forepaw, and removal behaviors were recorded for 3 min. **B)** Barplot showing number of removal behaviors (mean ± SEM) after injection of Ringer (solid bars) or lidocaine (hatched bars). Grooming (purple), licking (orange) and sticky behaviors (magenta; paw shaking, swiping, tapping pooled). Data were tested with a multiple comparisons ANOVA test (p = 0.0087 for treatment Ringer vs Lidocaine factor). Multiple comparisons Tukey’s test: adjusted p-values of p = 0.383 (grooming Ringer vs Lidocaine), p = 0.6019 (licking Ringer vs. Lidocaine) and p = 0.0017 (sticky Ringer vs. Lidocaine). Bar plots are mean ± SEM. ** p<0.01.

To examine whether forepaw anaesthesia altered the timing of removal behaviors rather than only their overall occurrence, we analysed the temporal distribution of individual behavioral events following Ringer and lidocaine injection (**Figure S2A**). For each rat and behavioral category, we quantified the half-rise time—the point at which 50% of that animal’s total events had occurred—as a measure of response latency (**Figure S2B**). Half-rise times did not differ between Ringer- and lidocaine-injected animals for any behavioral category (two-way ANOVA, main effect of treatment p = 0.196), indicating that lidocaine reduced the number of sticky behaviors without shifting the timing of the behavioral response relative to control injected animals.

### Somatosensory cortical neuron responses and differentiation with sticky and non-sticky stimuli

Having confirmed the role of forepaw afferents in the behavioral response to stickiness in pups, we next wondered about the cortical cellular responses underlying the perception of stickiness which might mediate such responses. Due to sparse firing responses and subthreshold encoding of touch in the somatosensory cortex ^9^, we performed patch-clamping *in vivo* to examine subthreshold and firing responses to sticky and non-sticky stimuli. We targeted recordings to the forepaw region of the somatosensory cortex ^10^. Once a whole cell recording was established, the sticky and non-sticky delivery apparatus was triggered, revealing membrane potential modulation with forepaw touch (**Figure 5A**). To understand how the somatosensory cortex may represent stickiness, individual neurons were assessed for (i) changes in firing rate and subthreshold responses at the onset and offset of the stimulus (ii) differential responses between stimuli and (iii) response categories across the population of analyzed neurons.

**Figure 5:**
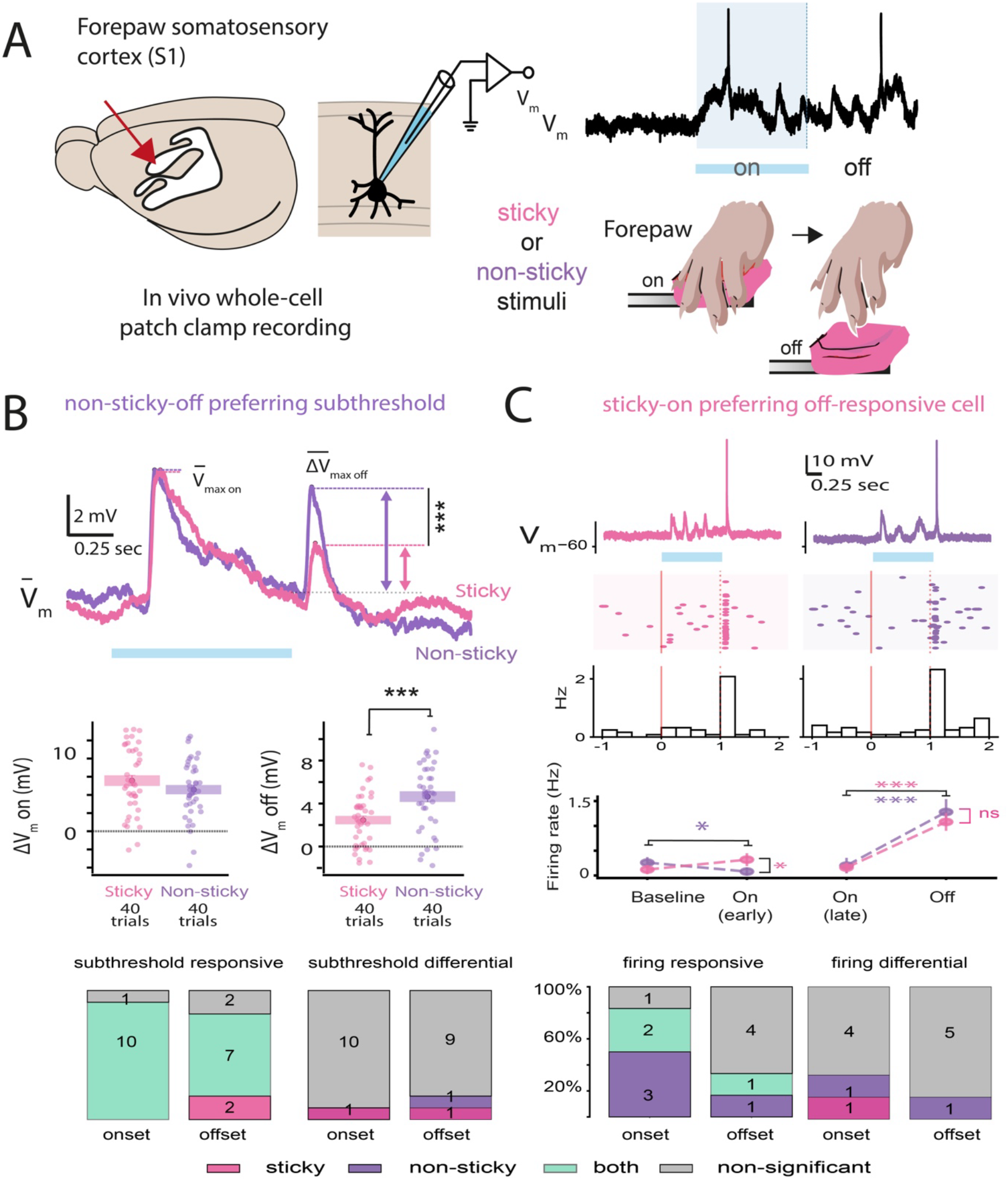
Responses and sparse differential preferences of somatosensory cortical neurons to sticky and non-sticky stimuli. **A)** Schematic indicating targeted location for whole-cell patch-clamping *in vivo* of neurons in the forepaw somatosensory cortex, during presentation of a sticky or non-sticky stimulus to the forepaw of the animal. **B)** (Top) Example of a non-sticky-off preferring subthreshold cell. This cell was responsive for sticky and non-sticky stimuli in the onset and offset periods. When comparing the change in V_m_ for onset sticky vs non-sticky over all trials (n = 40 sticky and n = 40 non-sticky), there was not a difference (p = 0.504, Mann Whitney U). When comparing the change in V_m_ for offset sticky vs non-sticky over all trials (n = 40 sticky and n = 40 non-sticky), there was a significant difference (p = 0.0003, Mann Whitney U; Median change in V_m_ sticky = 2.41 mV; Median change in V_m_ non-sticky = 4.98 mV). (Bottom) Barplots showing the proportion of cells, with significant subthreshold response to stimulus onset or offset and differential firing preferences. **C)** (Top) Example of an offset-responsive, onset differential firing neuron. Representative single-trial V_m_ traces of each neuron to non-sticky and sticky stimuli. Blue bars denote stimulus presentation period (1s). (Middle) Raster plots showing spike timings per trial and mean spike rates (binned by 0.25 s), aligned to stimulus onset. (Bottom) The firing rate response at stimulus onset from baseline is significantly different between sticky vs non-sticky (p = 0.004, Mann Whitney U, n=50 trials sticky, n=50 trials non-sticky) though not for offset (p=0.881, Mann Whitney U, n=50 trials sticky, n=50 trials non-sticky). Firing rate at stimulus offset responds significantly compared to pre-offset for sticky and non-sticky stimuli (p < 0.0001 and p < 0.0001 respectively, LMM). (Bottom) Barplots showing the proportion of cells, with significant firing rate response to stimulus onset or offset and differential firing preferences. Details of statistical testing are outlined in the methods section. * p < 0.05, ** p < 0.01, **** p < 0.0001, ns, no significant difference.

With sticky and non-sticky stimulus applications over trials, we observed subthreshold depolarizations following the stimulus onset as well as a subthreshold offset response, once the stimulus was removed, which was apparent in the average membrane potential (V_m_) across trials. In the example shown, we assessed the trial-by-trial V_m_, we confirmed responsiveness of the neuron to stimulus onset and offset. We additionally observed a differential subthreshold offset response, with non-sticky preference (n= 40 trials sticky and 40 trials non-sticky, **Figure 5B top and Figure S3A-B**). Cells were responsive to sticky and non-sticky stimuli at onset (91% subthreshold onset responsive for both) and offset (64% offset responsive for both sticky and non-sticky, 18% offset responsive to sticky). Differentially responding cells were sparse, with 9% showing onset differentiation and 18% showing offset differentiation (total 3 differential cells, **Figure 5B, bottom**).

Neurons which fired in more than 50% of trials were analyzed for firing responses. We display an example of a neuron which decreased its firing at onset to non-sticky stimuli, and was differential to the sticky onset. At offset, the neuron increased firing significantly for sticky and non-sticky stimuli, without differentiation (**Figure 5C, top**). Overall, firing cells were more responsive at onset, with 33% onset responsive for both stimuli and 50% responsive for non-sticky stimuli. 17% of cells were offset responsive for non-sticky and 17% offset responsive for both. Differentially responding cells were also few for firing analysis, with 33% showing onset differentiation (1 sticky and 1 non-sticky) and 17% (1 cell) showing offset differentiation with preference for non-sticky (**Figure 5B, bottom**).

Lastly, we analysed the depth and response profiles for neurons, and found that neurons overlapped in firing and subthreshold responsiveness, though differential firing and subthreshold cells were separate in identity and depth (**Figure S4**). Overall, we report mixed responsiveness of forepaw cortical neurons for sticky and non-sticky stimuli and sparse sticky vs non-sticky differential subthreshold and supra-threshold coding displayed in onset and offset periods of stimulus presentation.

## Discussion

Using quantitative behavioral analysis with sticky stimuli, local blockade of peripheral sensing and *in vivo* patch clamp electrophysiology, we characterized stickiness responses in rats. Stickiness evokes immediate and intense sticky-specific behaviors that are driven by forepaw afferents and, unlike responses to other tactile stimuli, are possibly related to stimulus detachment.

### Body part specific, intense and distinct stickiness responses in rats

A remarkable characteristic of the rat stickiness responses observed here is an unexpected degree of specificity. We expected rats to react to the application of adhesives to their body, but we were surprised by the extent of body axis-specificity we observed. One might expect responses to stimuli applied to the fur (e.g., the nape) to differ from those at the glabrous paw skin. Less expected was the degree to which responses differed between forepaw and hindpaw, two anatomically similar glabrous surfaces. We reason that the forepaw, which is used to engage in complex ways with objects (e.g., explore and manipulate) in rodents ^11,12^ and other mammals, harbor specific response mechanisms to stickiness.

We observed intense, fast onset behavioral responses to sticky stimuli that we (MB and AC personal observations) have not observed in decades of work with rats. Indeed, the response time course of sticky-specific was significantly different from grooming or licking behaviors. This finding suggests that sticky-specific behaviors differ from routine body care behaviors such as grooming^13^.

In line with our other observations, we found sticky-specific behaviors to be selective for the property of stickiness itself, rather than for any single stimulus. Accordingly, mochi and marshmallow—two sticky substances that differ in chemical composition, scent and taste—evoked similar behavioral responses that were distinct from reactions to water or oil stimuli. We suggest that stickiness defines the observed rat behavioral responses.

### Stimulus detachment, biophysics and neural mechanisms of stickiness

Several lines of evidence point to the significance of stimulus detachment in the emergence of stickiness sensation. In the behavioral domain, we wonder if the paw tapping behavior we observed in rats is particularly informative for enabling rapid sampling of stimulus detachments. Tapping is very different from the more vigorous swiping and shaking behaviors and it seems well suited to enable the sensing of stickiness by repeated stimulus detachments.

In the human literature, there are multiple leads pointing to the significance of stimulus detachment for stickiness perception. The work of Nam et al. 2020 ^2^ related forces resulting from stimulus detachment to stickiness judgements in humans. Similarly, in experiments, in which liquid viscosity was manually experienced, Cavdan et al. identified pull-off forces as related to unpleasant and possibly sticky percepts ^4^. Detailed analysis of whisking on sticky surfaces indicates stick-slip events and rolls as defining metrics in active exploration of stickiness ^3^, which could provide a whisker analogue of our observed forepaw detachment-mediated sticky behaviours.

Our work not only showed that rat forepaws are highly sensitive to stickiness, but we also demonstrated that forepaw afferents are decisive drivers of this sensation. Specifically, we observed a loss of stickiness responses following afferent blockade by lidocaine injections.

Moving from peripheral sensors of stickiness to cortical representations, we then report, for the first time to our knowledge, differential single-cell responses to stickiness in the somatosensory cortex. The evidence of such cells, in naive animals no less, implies an intrinsic dimension by which the cortex and its downstream projections may distinguish stickiness, enabling the instantaneous perception of stickiness via the forepaw that could elicit the sticky-specific behaviors reported.

Additionally, the specificity of some differential responses to the onset and offset periods further implies distinct perceptual encoding of stimulus attachment and detachment. This could be especially revealing, given that behaviors such as tapping may constitute sampling of sticky-specific sensory features at attachment and detachment. These responsive neurons thus provide the building blocks needed for a specific encoding of stickiness in the cortex, sensed peripherally via afferents in the forepaw.

The diversity of responses we measured in the somatosensory cortex could imply the simultaneous and heterogeneous encoding of multiple sensory features, which supports higher-dimensional representations and complexity ^14^. Stickiness is a complex percept, likely relating to multiple sensory features including touch, texture, and more. Further investigations into the neural representation of stickiness could capitalize on high-density electrophysiological methods such as Neuropixel recordings to record spiking activity from many neurons at once, potentially during awake behavior and with more stimulus types spanning a broader range of stickiness, to further understand the nature and geometry of stickiness representation in the brain.

Furthermore, the richness and distinct cellular encoding of subthreshold responses to stickiness should be progressed in parallel. The finding that separate neurons differentiate stickiness and non-stickiness with firing and subthreshold responses indicates distinct intrinsic properties and potentially cortical neuron cell-types. Whether and how subthreshold and suprathreshold sticky-encoding neurons interact across cortical layers may be a further step towards understanding the circuit-level computation of the sticky percept.

### Conclusions: a stickiness response system

We observed a distinct set of stickiness-related behaviors in response to sticky stimuli. Such behaviors were not only specific to sticky stimuli, but also occurred at shorter latencies than grooming or licking behaviors. It appears that shaking and swiping behaviors aim at stickiness removal. More generally we would place these vigorous short-latency behaviors in the defensive category. This behavioral categorization would fit with human data that indicate aversiveness of sticky sensations ^4,15^. We assume that these behaviors likely are part of an innate stickiness response system. All pups from different litters showed such behaviors. Given their young age and sensorially restricted upbringing in a rat cage it is likely that these animals never experienced stickiness before, yet they showed highly consistent behaviors.

We also wonder if stickiness responses are evolutionarily old; informal observations on humans indicate that they also evaluate stickiness by detaching their fingers and they also seem to be keen to remove sticky stimuli from their hands. Together, these findings position stickiness as a distinct somatosensory dimension with its own behavioral and neural signatures. Despite its ubiquity in everyday tactile experience, stickiness has received little attention in systems neuroscience, and human stickiness responses and their potential continuity with rat behaviors deserve further investigation.

## Methods

### Animals

Male and female Long evans wild-type rats were used in this study (ages P21 to P31). All procedures were approved by the local ethics committee at the Marine Biological Laboratory in Woods Hole (IACUC, protocol 26-09A).

### Behavioral experiments

Experiments were conducted in an arena and filmed from a side and top view (Logitech). Video acquisition was conducted using OBS Studio v32.1.2 (Open Broadcaster Software). All stimuli were delivered by syringe, delivering 0.1 mL to the specified location immediately before placing animals in the behavior arena. Marshmallow stimuli consisted of Marshmallow fluff (Durkee-Mower, Inc., Lynn, MA). The mochi stimulus (Koda Farms mochiko sweet rice flour) was equal parts mochiko flour and water, mixed, microwaved for 30 seconds and cooled to room temperature. The oil stimulus was sunflower seed oil (Ahold, USA). The water stimulus was tap water from the Marine Biological Laboratory faucet.

### Behavioral scoring and analysis

Five behaviors were logged: grooming, shaking, tapping, swiping, and licking. Grooming was defined as the stereotyped syntactic grooming chain^16^: an ordered progression from rapid small ‘ellipse’ forepaw strokes over the nose and vibrissae, through unilateral and then large bilateral strokes across the face and head. Forepaw shaking was defined as rapid oscillatory movements of an elevated forepaw, or of both forepaws together. Forepaw swiping was defined as alternating sweeps of the left and right forepaws across the floor. Forepaw tapping was defined as brief, repeated lifting and replacement of the forepaws against the floor. Licking was defined as licking any part of the body or adhesive, whether as part of self-grooming or not.

After application of 0.1 mL stimuli, the rat was placed in the ‘rat-a-viewey’ observation chamber. This consisted of a transparent box with two mounted cameras, one above and one beside the chamber to record behavior from multiple viewpoints simultaneously. Behaviours were scored live during each trial using a custom programme and graphical user interface written in Python (‘Rat-a-GUI’). A trial was initiated via ‘Rat-a-GUI’ as soon as the rat was placed in the ‘rat-a-viewey’ observation chamber, with occasional delay of up to 15s due to device lag. During a trial, the experimenter pressed a button, or the matching number key, each time the animal performed one of the five scored behaviours. The application recorded the time of each behavior as seconds elapsed from trial onset. Each trial ran for a fixed 180 second window after the start of ‘Rat-a-GUI’ trial initiation, after which scoring stopped automatically. On completion the application appended a single row per trial to a spreadsheet, storing every event time and the per-behaviour event count together with the animal ID, adhesive, body location, and trial duration.

For removal analysis in Figure 1, If the adhesive was not successfully removed within the 600s trial, the proportion of time was assigned a ceiling value of 1.0. Trials in which removal occurred without an observable foregrooming response were excluded from analysis rather than assigned a ceiling value, as this reflects rapid removal instead of no response.

To assess the anterior to posterior relationship of behaviors, categories of body parts were assigned into nose, dorsal/ventral forepaw and dorsal/ventral hindpaw, in anterior-posterior ordering. Nape was not included because it was unclear where this location fell on the relative axes. A mixed-effects linear trend analysis was run to determine whether the slope of behavioral measures across animals were significantly non-zero.

### Blocking of afferents

To block tactile sensation, 0.05 mL of 0.5% lidocaine diluted in Ringer solution were injected to the dorsal forepaw, controls were injected with 0.05 mL of Ringer solution. After 5 minutes, 0.1 mL of marshmallow was applied to the dorsal forepaw and behavior was recorded for 3 minutes, scoring five behaviors (grooming, shaking, tapping, swiping, and licking) as described previously.

### *In vivo* electrophysiology

Whole-cell patch-clamping was performed *in vivo* in rat pups under terminal anaesthesia (urethane 1.4 g/kg, administered intraperitoneally). Loss of consciousness was confirmed by toe-pinch tests before surgery was performed. Briefly: 1% lidocaine was injected under the scalp before incision. Animals were head-fixed on a stereotaxic surgery stage and kept warm. Body temperatures were monitored via a rectal thermal probe and maintained at approximately 32–33 C. An incision was placed in the scalp to expose the skull, and a craniotomy was made using a dental drill over the forepaw somatosensory cortex^10^ (approximately -0.5 mm anterior and +3.5 mm lateral of Bregma).

Patch electrodes were pulled from borosilicate glass with filament (Sutter Instruments glass) on a Model P-1000 Micropipette puller (Sutter Instruments), with an estimated resistance of 3–5 MOhm. Pipettes were filled with sterile-filtered intracellular saline. Recordings were performed largely following published methods^17–19^. Membrane potential was recorded at a sample rate of 25 kHz in current clamp mode. At the end of each recording, the depth of the cell was estimated from the distance that the micromanipulator had lowered into the brain from the zeroed cortical surface.

### Electrophysiology stimulus delivery

A custom stimulus apparatus was constructed in which a lever pivoted about a central fulcrum, mounted on a micromanipulator (Narishige), to deliver a stimulus to the ventral forepaw from below. An airpuff directed at the distal end of the lever pushed it downward, raising the proximal end—which carried the stimulus—upward into contact with the forepaw. The forepaw was gently rested on a beam and secured with clay to ensure stereotyped, reproducible stimulation. The airpuff was triggered under control of the custom Spike2 (CED Instruments) protocol and directed at the lever from a distance of approximately 25 cm from the animal to avoid whisker deflection.

A piece of sticky thermoplastic rubber (PICcircuit via Amazon) was used as a sticky stimulus, the same rubber covered with plastic film served as a non-sticky control. Cell responses were recorded during trains of 10 stimulations, each lasting 1 s with a 5 s inter-stimulus interval.

### Electrophysiology analyses

The recorded membrane potential (V_m_) from each cell was aligned to the onset of each sticky or non-sticky stimulation trial (duration 1-s), and separated into baseline (1-s pre-onset), stimulation, and post-offset epochs for analysis. To ensure that only trials with healthy resting membrane potentials (RMPs) were analyzed per cell, the RMP of each trial was calculated by taking the mean and median V_m_ of each trial baseline 0.5 seconds before the stimulus onsets. Trials in which the RMP was higher than -30 mV were excluded. Only cells with at least 10 such quality-controlled trials per sticky and non-sticky stimulus type recorded were included in downstream analyses.

To further ensure that analysis included cells with physiological responses above noise, we calculated a signal-to-noise measure consisting of the mean onset membrane potential response divided by the median standard deviation of the baseline 50 ms window. Cells with a signal-to-noise ratio below 2 were not included in subthreshold analyses. These criteria were applied to 23 recorded cells, resulting in 11 cells analysed as described below for subthreshold responses and 6 cells for firing rate responses. Two firing rate analysis cells were not analyzed for subthreshold responses due to the signal-to-noise criteria.

### Firing neuron classification and analyses

A cell was classified as a firing neuron if it spiked in more than 50% of stimulation trials. Action potential detection was performed by first filtering the voltage trace using a lowpass filter (scipy.signal.butter) with a cutoff of 2000 Hz, and then calculating the rate of voltage change (dV/dt) in mV/ms. Spike times were then identified by the time of each dV/dt peak exceeding 10 V/ms (scipy.signal.find_peaks).

The firing rate of each neuron to the baseline, stimulation onset, pre-stimulation offset and stimulation offset epochs of each trial were calculated by taking the mean firing rate over 1 s pre-onset, 0.5 s post-onset, 0.5 s pre-offset and 0.5 s post-offset respectively. Post-onset and -offset responses were restricted to the first 0.5 s to limit analysis to temporally-locked responses.

To classify if a cell was responsive to stimulation in the onset or offset and whether they were differential, paired tests (parametric or non-parametric, following tests for normality) were conducted to compare the trial by trial firing rate of a neuron 0.5 s after onset or offset to its firing rate at pre-stimulus baseline or 0.5 s pre-offset respectively (alpha = 0.05). Neurons in which the change in onset/offset across trials were judged to be different between stimuli were classified differential. To provide a second confirmation of whether a cell responded to stimuli, a linear mixed effect model was used to compare the firing rate of each neuron in a trial before and after stimulation onset / offset with either stimulus type (sticky versus non-sticky), grouping by trial to account for trial-to-trial differences (statsmodels.formula.api.mixedlm). All paired tests of firing rate agreed with significance categories follow the LMEM analysis. The preferred stimulus type of a differential neuron was that for which the absolute magnitude of its firing rate change was greater.

### Subthreshold analyses

The average V_m_ trace was calculated across trials for sticky and non-sticky and V_m_ max was measured within the stimulus period as well as from the offset to 1s after offset (depicted in **Figure S2** for examples). The time of V_m_ max (in stim) and V_m_ max (post stim) was then used to measure V_m_ across trials at a set time. To classify a cell as responsive or not responsive to each stimuli onset, the paired V_m_ at T_0_ (stim onset) was compared with V_m_ max (in stim) and statistically tested over trials. For responsive classification at offset, a paired comparison between V_m_ at T offset and V_m_ at V_m_ max avg (post stim). Delta V onset, was calculated as V_m_ max (in stim) minus V_m_ at T_0_; Delta V offset, was calculated as V_m_ max (post stim) minus V_m_ at T offset for every trial for sticky and non-sticky.

## Data and code availability

All underlying data, media and code is deposited and will be made publicly available upon publication.

## Associated videos 1–4, also described in results section

Video 1: Sticky shaking video: https://figshare.com/s/9f762fa75db9e7c34801

Video 2: Sticky swiping video: https://figshare.com/s/2dc702f9f5728320e62f

Video 3: Sticky taps video: https://figshare.com/s/8d25b4aebcde6f11a2a2

Video 4: Sticky behaviors all video: https://figshare.com/s/47b7b4370518222740eb

## Supplementary Materials

**Videos 1-4 described in results and supporting Figure 2**:

**Video 1: Sticky shaking video**: https://figshare.com/s/9f762fa75db9e7c34801

**Video 2: Sticky swiping video**: https://figshare.com/s/2dc702f9f5728320e62f

**Video 3: Sticky taps video**: https://figshare.com/s/8d25b4aebcde6f11a2a2

**Video 4: Sticky behaviors all video**: https://figshare.com/s/47b7b4370518222740eb

---

**Supplementary Figure 1:**
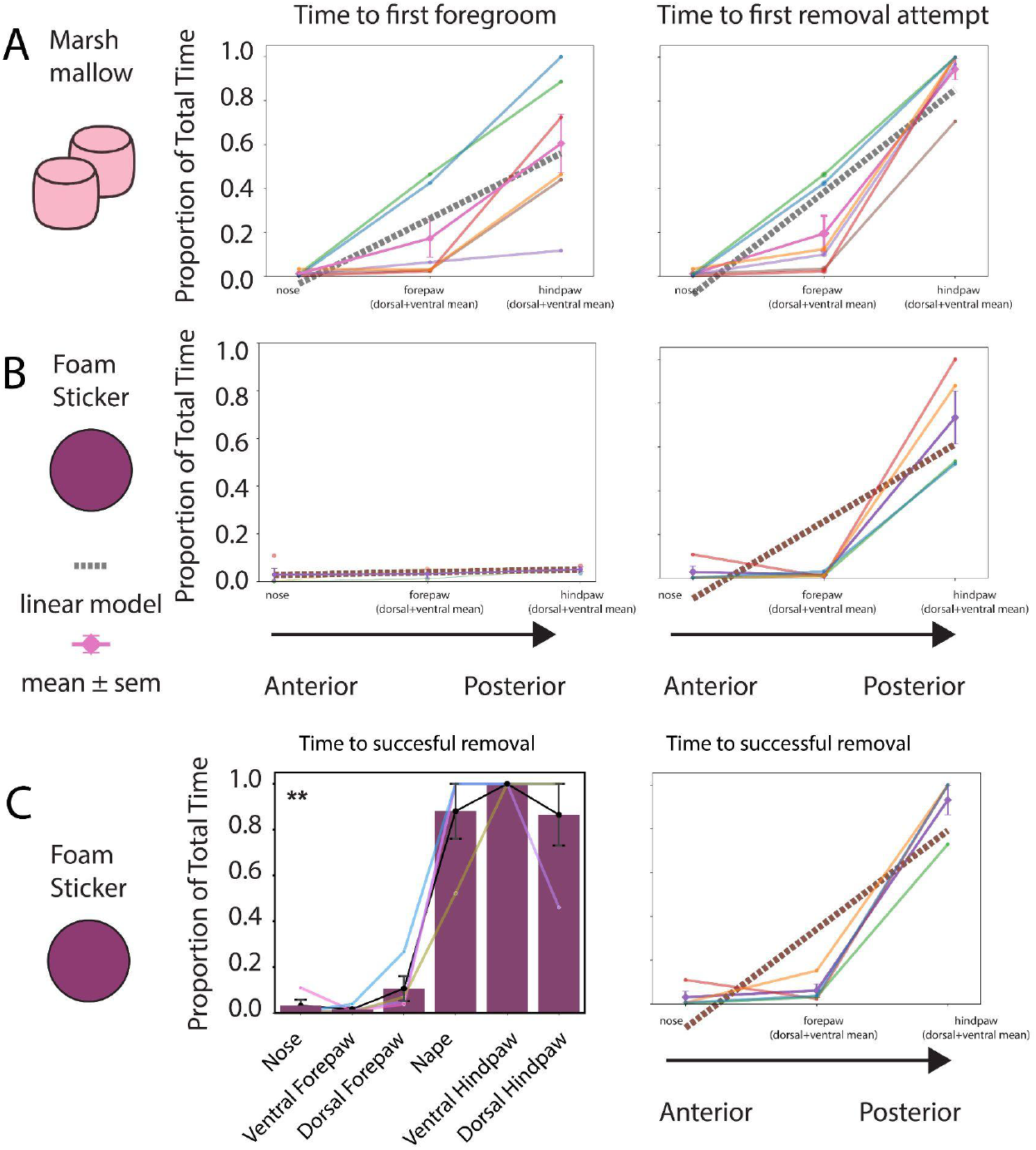
Anterior to posterior analysis of behavioural measures in response to sticky stimulus application across body parts. Anterior-to-posterior behavioral responses were analyzed based on nose (most anterior), the mean of dorsal and ventral forepaw response and the mean of dorsal and ventral hindpaw response (most posterior). **A)** Data for marshmallow stimulus: A mixed effects linear analysis was performed on behavioral data for 6 animals (dotted line). Mean and SEM are depicted as pink lines with error bars. Each colored line represents data for an animal. As in Figure 1, metrics are First foregroom attempt (left) and first removal attempt (right). **B)** Data for foam sticker stimulus: same conventions as in A. p-values are reported in Figure 1. **C)** Left: Barplot showing the proportion of time taken to successful removal of foam sticker across body locations (Friedman test, Q = 17.81, p = 0.0032). Right: mixed effects linear analysis performed on time taken to successful removal for 6 animals (dotted line, p = 2.30e-09). ** p < 0.01

**Supplementary Figure 2:**
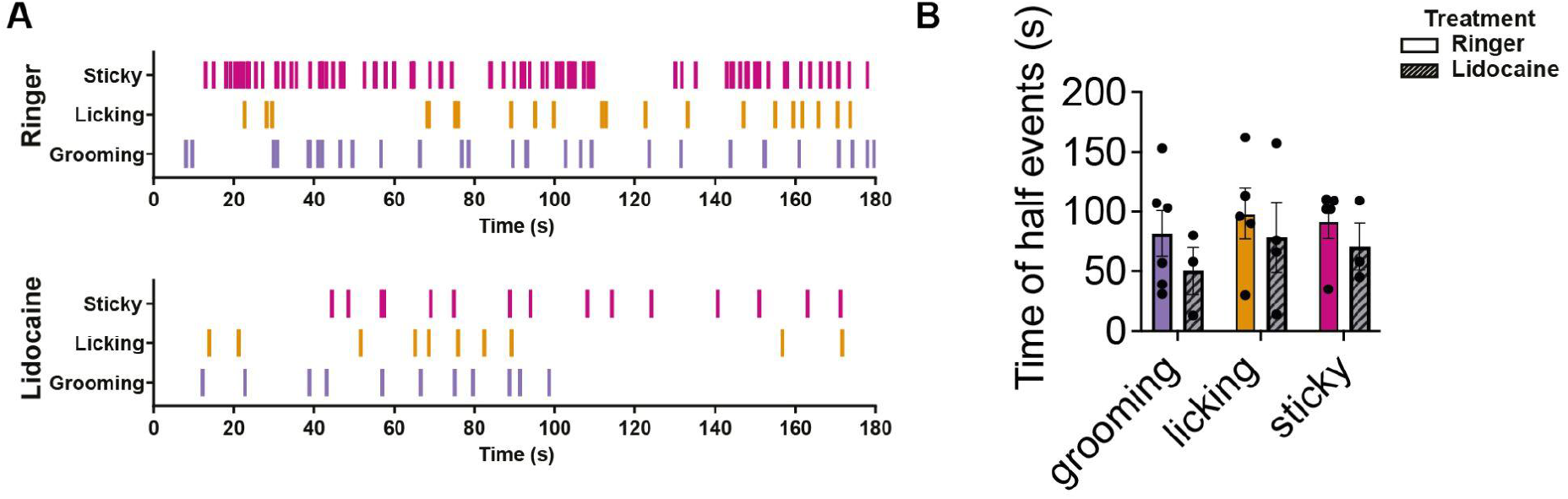
Temporal characteristics of behavioral responses following local Ringer and lidocaine injections. **A)** Raster plots showing individual behavioral events, categories are the same as in Figure 4B. Events are summed across all rats. **B)** Half-rise times, defined for each rat and behavioral category as the time when 50% of total observed events were performed. Animals with no observed events of a category were excluded from that category. No statistical differences were observed when comparing Ringer vs lidocaine injected animals. Data were tested with a multiple comparisons ANOVA test (p=0.196 for treatment Ringer vs Lidocaine factor). N=5 rat lidocaine, N=6 rats Ringer. Bar plots are mean ± SEM.

**Supplementary Figure 3:**
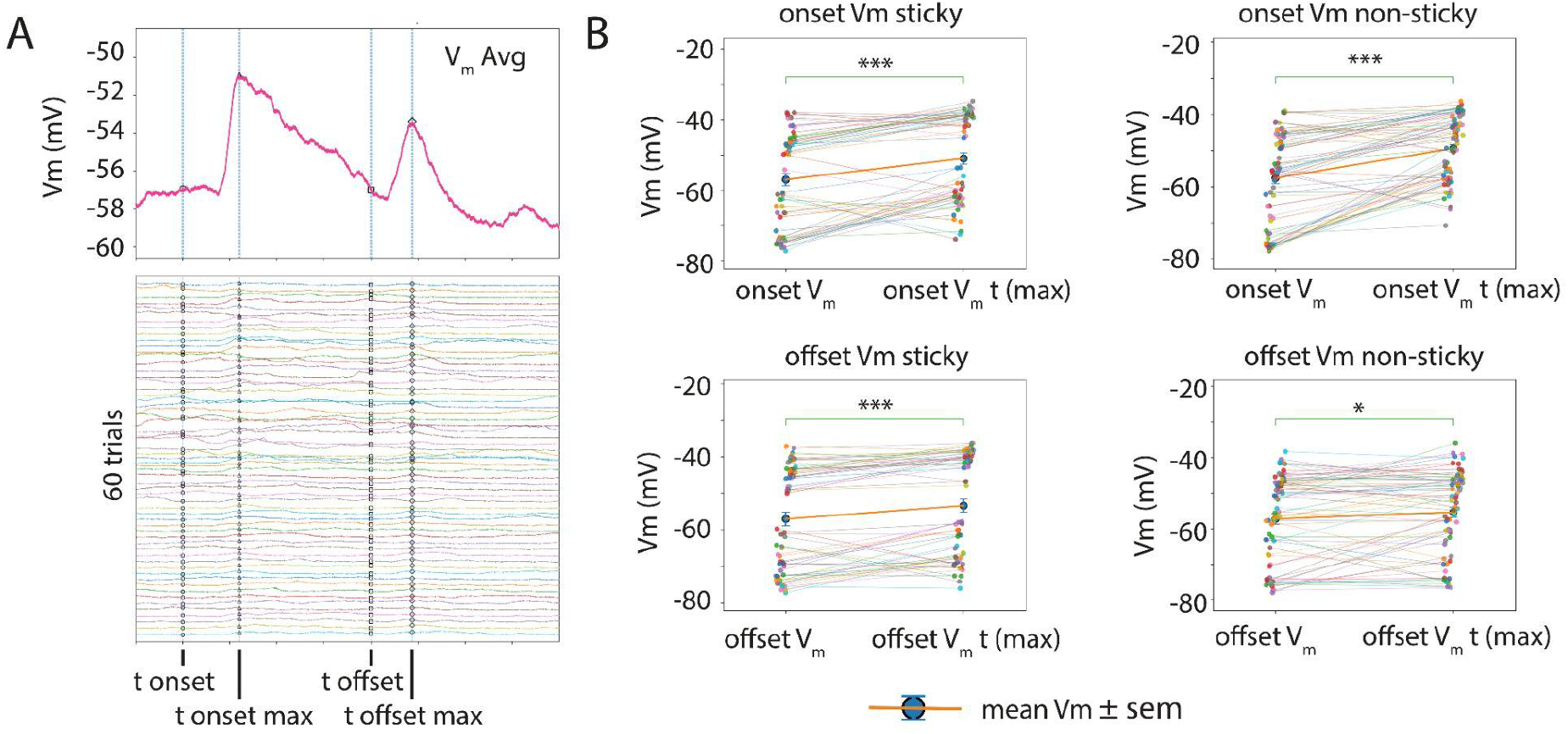
Steps of analysis for data in Figure 5. **A)** Mean traces are calculated for each stimuli and used to determine the V_m_ onset time and offset time across trials. Across trials, the Vm at onset, onset max, offset, and offset max were taken as the mean of 10 ms, 50 ms, 10 ms and 50 ms windows, respectively. **B)** Trial based measures were then compared across all trials and tested using paired statistics as well as a linear mixed effects model (LMM). If significance was found for both tests, the cell was considered responsive for the given stimuli. This cell responded significantly for subthreshold onset and offset for both sticky and non-sticky stimuli. * p < 0.05, *** p < 0.0001

**Supplementary Figure 4:**
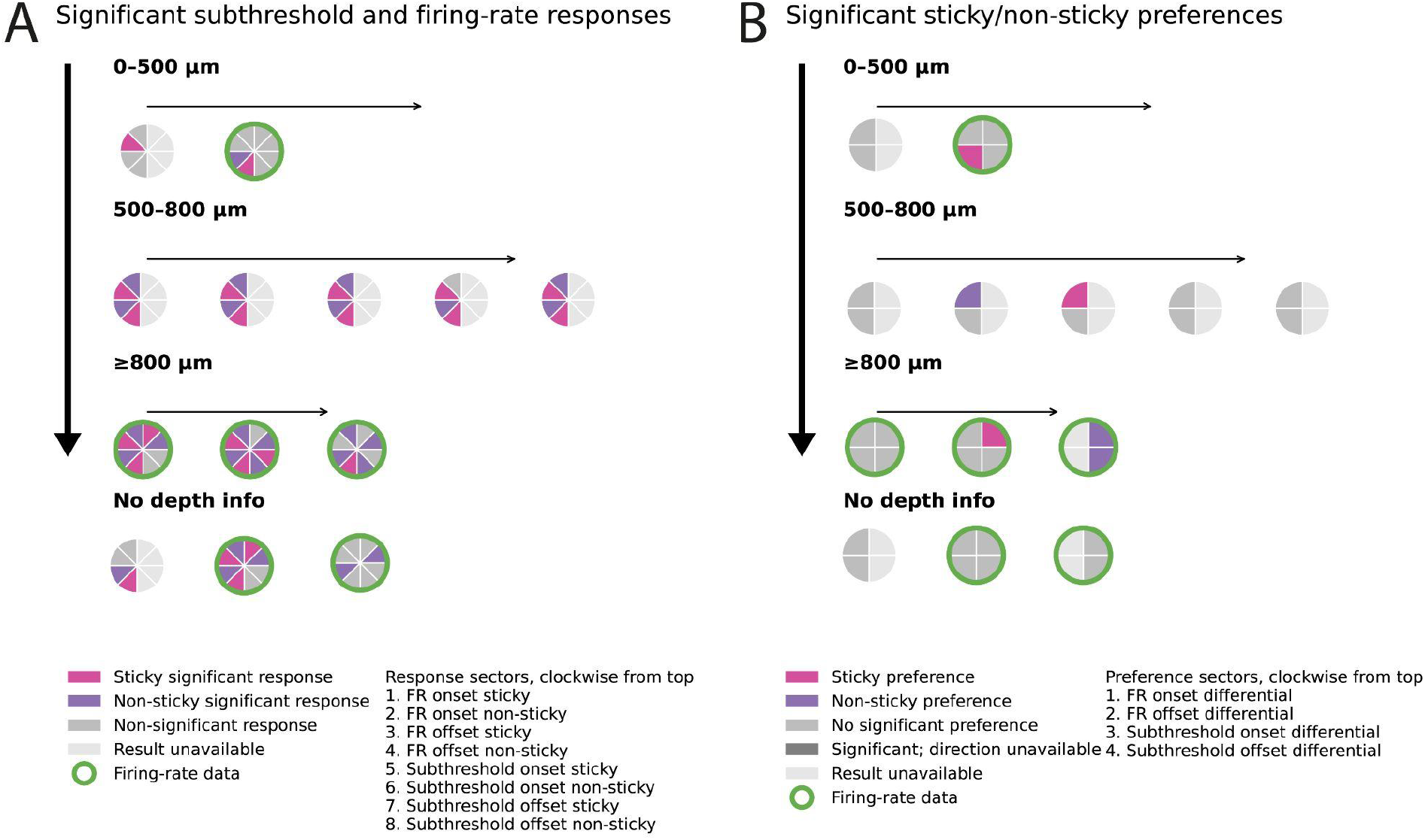
Responses and differential cells by identity and cortical depth. **A)** Individual cell profiles for all neurons analyzed for onset and offset significance across depth (cells without depth info also shown). **B)** Individual cell profiles for all neurons analyzed for differential sticky vs non-sticky responses across depth (cells without depth info also shown). Significance was classified with alpha <0.05.

## Acknowledgements

We thank Luigs & Neumann, NPI electronic, Cambridge Electronic Design (CED) and other vendors for loan equipment, the Neural Systems and Behavior course at the Marine Biological Laboratory, the forever people Bruce Carlson, Lauren O’Connell, Rosalie Maltby, the fish, fly modules and the 2026 NS&B class for collegial support.

## References

1. Hanada, M. (2020). Food-texture dimensions expressed by Japanese onomatopoeic words. Journal of Texture Studies 51, 398–411. 10.1111/jtxs.12499.

2. Nam, S., Vardar, Y., Gueorguiev, D., and Kuchenbecker, K.J. (2020). Physical Variables Underlying Tactile Stickiness During Fingerpad Detachment. Front. Neurosci. 14, 235. 10.3389/fnins.2020.00235.

3. Wyche, I.S., O’Neil, M.A., and O’Connor, D.H. (2026). Whisker-based pre-neuronal and peripheral encoding of surface stickiness. Preprint, 10.7554/eLife.111939.1 10.7554/eLife.111939.1.

4. Cavdan, M., Fehlberg, M., Bennewitz, R., and Drewing, K. (2025). Gooey stuff: the psychophysics of unpleasantness in response to touching liquids. Proc. R. Soc. B. 292, 20252244. 10.1098/rspb.2025.2244.

5. Saluja, S., and Stevenson, R.J. (2022). Tactile disgust: Post-contact can be more disgusting than contact. Quarterly Journal of Experimental Psychology 75, 652–665. 10.1177/17470218211043688.

6. Kim, J., Yeon, J., Ryu, J., Park, J.-Y., Chung, S.-C., and Kim, S.-P. (2017). Neural Activity Patterns in the Human Brain Reflect Tactile Stickiness Perception. Front. Hum. Neurosci. 11, 445. 10.3389/fnhum.2017.00445.

7. Yeon, J., Kim, J., Ryu, J., Park, J.-Y., Chung, S.-C., and Kim, S.-P. (2017). Human Brain Activity Related to the Tactile Perception of Stickiness. Front. Hum. Neurosci. 11. 10.3389/fnhum.2017.00008.

8. So, Y., Kim, S.-P., and Kim, J. (2020). Perception of surface stickiness in different sensory modalities: an functional MRI study. NeuroReport 31, 411–415. 10.1097/WNR.0000000000001419.

9. Crochet, S., Poulet, J.F.A., Kremer, Y., and Petersen, C.C.H. (2011). Synaptic Mechanisms Underlying Sparse Coding of Active Touch. Neuron 69, 1160–1175. 10.1016/j.neuron.2011.02.022.

10. Chapin, J.K., and Lin, C. -S (1984). Mapping the body representation in the SI cortex of anesthetized and awake rats. Journal of Comparative Neurology 229, 199–213. 10.1002/cne.902290206.

11. Ruder, L., Schina, R., Kanodia, H., Valencia-Garcia, S., Pivetta, C., and Arber, S. (2021). A functional map for diverse forelimb actions within brainstem circuitry. Nature 590, 445–450. 10.1038/s41586-020-03080-z.

12. Barrett, J.M., Raineri Tapies, M.G., and Shepherd, G.M.G. (2020). Manual dexterity of mice during food-handling involves the thumb and a set of fast basic movements. PLoS ONE 15, e0226774. 10.1371/journal.pone.0226774.

13. Bolles, R.C. (1960). Grooming behavior in the rat. Journal of Comparative and Physiological Psychology 53, 306–310. 10.1037/h0045421.

14. Nogueira, R., Rodgers, C.C., Bruno, R.M., and Fusi, S. (2023). The geometry of cortical representations of touch in rodents. Nat Neurosci 26, 239–250. 10.1038/s41593-022-01237-9.

15. Kawabe, T., Morisaki, T., and Ujitoko, Y. (2026). Pseudo-slimy: A novel phenomenon to evoke stickiness perception. i-Perception 17, 20416695261458511. 10.1177/20416695261458511.

16. Cromwell, H.C., and Berridge, K.C. (1996). Implementation of Action Sequences by a Neostriatal Site: A Lesion Mapping Study of Grooming Syntax. J. Neurosci. 16, 3444–3458. 10.1523/JNEUROSCI.16-10-03444.1996.

17. Margrie, T.W., Brecht, M., and Sakmann, B. (2002). In vivo, low-resistance, whole-cell recordings from neurons in the anaesthetized and awake mammalian brain. Pflugers Archiv European Journal of Physiology 444, 491–498. 10.1007/s00424-002-0831-z.

18. Clemens, A.M., Wang, H., and Brecht, M. (2020). The lateral septum mediates kinship behavior in the rat. Nature Communications 11, 3161. 10.1038/s41467-020-16489-x.

19. Clemens, A.M., Lenschow, C., Beed, P., Li, L., Sammons, R., Naumann, R.K., Wang, H., Schmitz, D., and Brecht, M. (2019). Estrus-Cycle Regulation of Cortical Inhibition. Current Biology 29, 605–615.e6. 10.1016/j.cub.2019.01.045.

